# CHAP: A versatile tool for the structural and functional annotation of ion channel pores

**DOI:** 10.1101/527275

**Authors:** Gianni Klesse, Shanlin Rao, Mark S.P. Sansom, Stephen J. Tucker

## Abstract

The control of ion channel permeation requires the modulation of energetic barriers or ‘gates’ within their pores. However, such barriers are often simply identified from the physical dimensions of the pore. Such approaches have worked well in the past, but there is now evidence that the unusual behaviour of water within narrow hydrophobic pores can produce an energetic barrier to permeation without requiring steric occlusion of the pathway. Many different ion channels have now been shown to exploit ‘hydrophobic gating’ to regulate ion flow, and it is clear that new tools are required for more accurate functional annotation of the increasing number of ion channel structures becoming available. We have previously shown how molecular dynamics simulations of water can be used as a proxy to predict hydrophobic gates, and we now present a new and highly versatile computational tool, the Channel Annotation Package (CHAP) that implements this methodology.

## Introduction

Ion channels form pores that facilitate the selective movement of ions across lipid bilayers. They therefore control a wide range of vital physiological functions and represent an important class of therapeutic targets. However, these pores are also highly dynamic structures that switch between functionally closed and open states in response to different physiological stimuli, a process referred to as gating ^1^.

Elucidating the molecular mechanisms that underlie these gating processes remains an important objective for ion channel structural biology and recent advances in techniques such as cryo-electron microscopy are now generating an increasing wealth of new ion channel structures, including many captured in different conformational states (**Figure 1**). However, before this structural information can be compared to the detailed electrophysiological data that can often be obtained for many ion channels, the conductive status of any new structure needs to be known. Conventionally, this has been achieved by determining the physical (i.e. steric) dimensions of the channel pore. A variety of approaches and biomolecular visualization tools exist for determining the dimensions of pores, tunnels and cavities in proteins^2^, but by far the most common software used for mapping transmembrane ion channel pores are programs such as CAVER^3,4^ and HOLE^5^. These calculate a radius profile along the conduction pathway and the structure is typically considered to be open and conductive if its narrowest constriction exceeds the radius of the hydrated ion species it conducts (**Figure 1A**).

**Figure 1:**
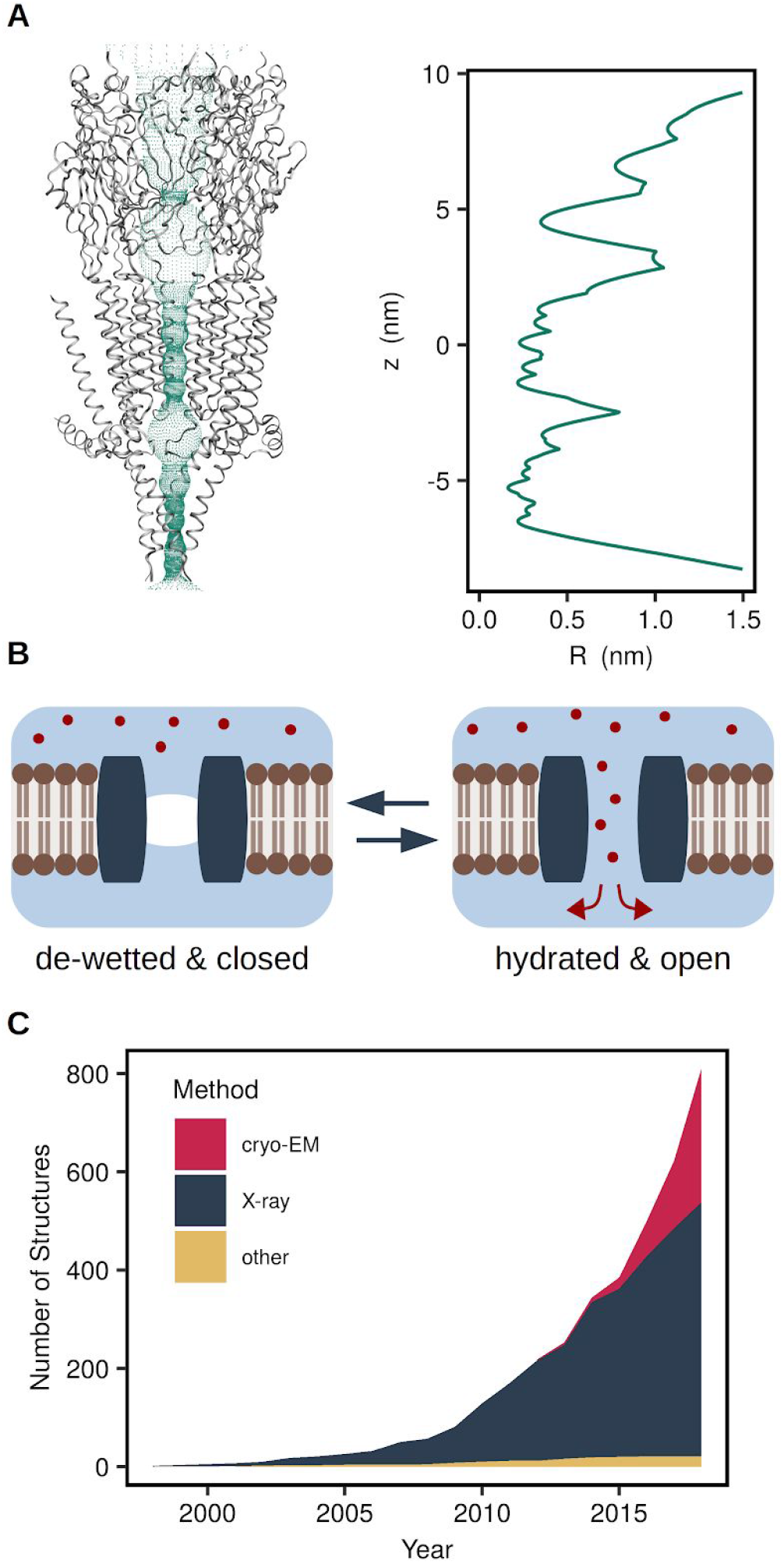
Hydrophobic gates pose a challenge to the functional annotation of ion channel structures. (A) HOLE profile of the entire PDB structure of a 5-HT_3_ receptor (PDB ID: 4PIR). (B) Hydrophobic gating: A non-conducting state is characterised by the de-wetting of a part of the permeation pathway lined by hydrophobic residues. Ions are prevented from crossing this region by the free energy barrier associated with their desolvation even if the pore is sufficiently large to accommodate them. (C) The number of ion channel structures deposited in the protein data bank is growing exponentially (data retrieved on 5 December 2018).

This size-based approach has served the ion channel community well for several decades, but an increasing body of evidence has demonstrated this approach is unable to detect gates that are not based on steric occlusion of the permeation pathway. For example, ‘hydrophobic gates’ can occur in channel pores to prevent ion flow and rely on the capillary evaporation of water that occurs when the lining of the pore is particularly hydrophobic ^6^ In these situations, ion movement through these ‘de-wetted’ sections of the pore is prevented by the free energy barrier associated with stripping the ions of their hydration shell (**Figure 1B**) rather than by van der Waals clashes with tightly packed pore residues ^7^. It is important to note that this definition of hydrophobic gating excludes cases where pore de-wetting immediately precedes the collapse and/or constriction of hydrophobic pore segments involved in steric closure due to a conformational change in the protein ^8^

The phenomenon of hydrophobic gating was first observed in simulations of simple model pores ^9–11^ and has been confirmed experimentally in nanopores^12^. It has been found to occur within pores up to 1.2 nm in diameter and so includes the dimensions of most ion channel pores ^13^ Therefore, an ion channel can be functionally closed even if the pore appears physically wide enough to accommodate the relevant hydrated ion.

The first example of a hydrophobic gate in an ion channel was found in an early cryo-EM structure of the nicotinic acetylcholine receptor, which represents a functionally closed state without being sterically occluded ^14^ Further examples of hydrophobic gates and barriers include the MscL mechanosensitive channel ^15^, the cation-selective bacterial channel GLIC ^16^, the BK and TWIK-1 potassium channels ^17,18^, the BEST1 chloride channel^19^, and the CorA magnesium channel ^20^ Thus hydrophobic gating clearly represents an important mechanism for regulating ion channel permeation and needs to be taken into consideration when assessing the functional status of any new ion channel structure.

In a previous study we have shown how molecular dynamics (MD) simulations of water behaviour within these nanoscale structures can be used to aid the functional annotation of different pentameric ligand-gated ion channels (the Serotonin and Glycine Receptors) and demonstrated the existence of hydrophobic gates within these particular structural conformations ^21^. In particular, we established that pore hydration observed in simple equilibrium MD simulations can be used as a reliable proxy for ion permeability. This was also validated by comparison to more computationally expensive and time-consuming approaches such as umbrella-sampling free energy calculations and computational electrophysiology simulations. Here we now present a new and highly versatile computational tool for use within the structural biology community that exploits the principle of pore hydration to rapidly predict the conductive status of new ion channel structures.

## Workflow for ‘Channotation’ of Pore Structures

The workflow underlying this Channel Annotation Package (CHAP) is shown in **Figure 2** and the required software is freely available (www.channotation.org). As an initial analysis step, CHAP can be run directly on the PDB structure of any channel or nanopore to generate both radius and hydrophobicity profiles of the putative ion conduction pathway. These initial steps are easily performed on an average desktop computer and do not require MD simulation of the structure. However, if these profiles suggest the channel may contain a hydrophobic gate or any other interesting features then a more in-depth analysis based on equilibrium MD simulation of the structure can be performed.

**Figure 2:**
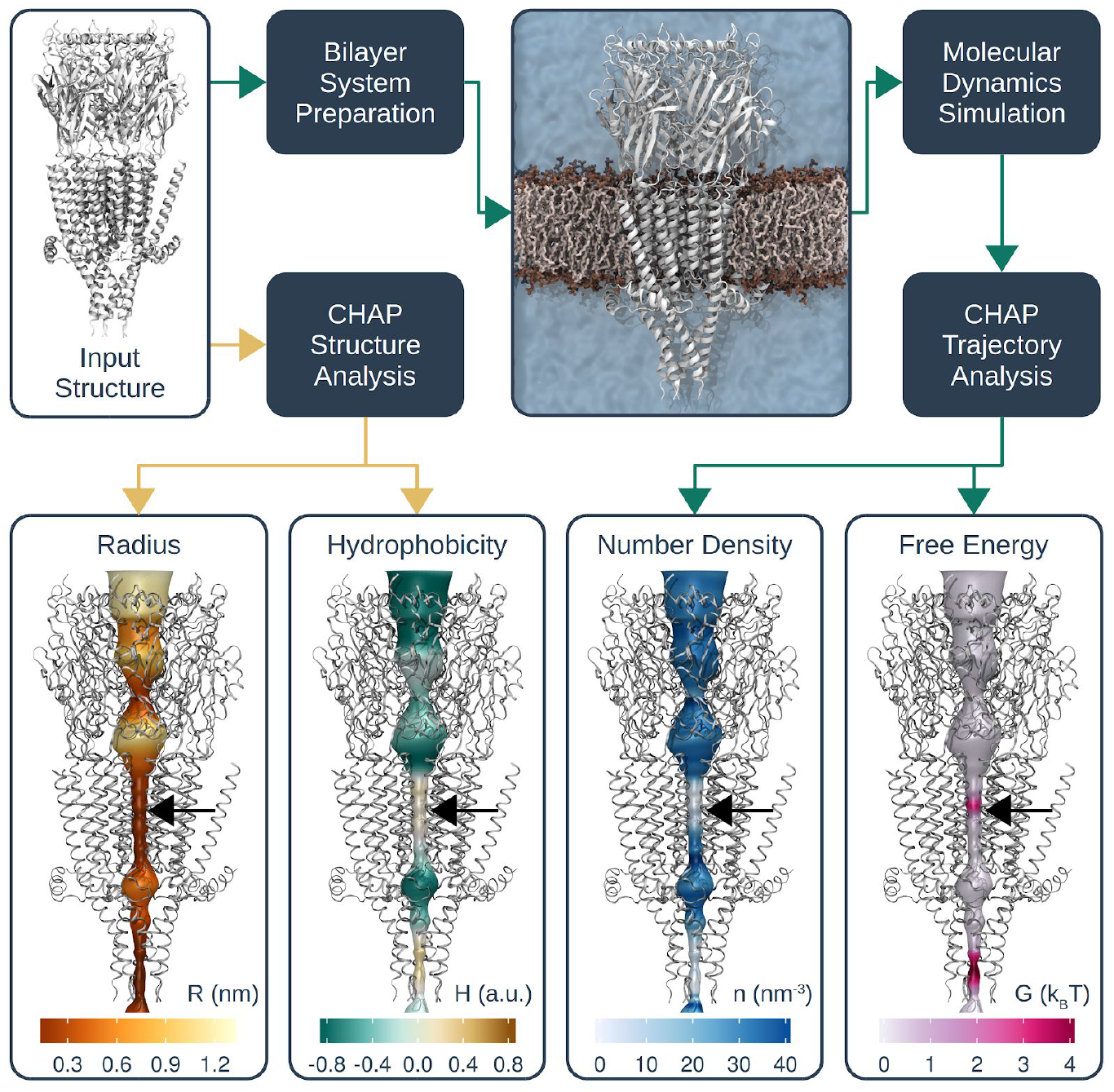
Overview of workflow for functional annotation of ion channels with CHAP. Given a simple protein structure as input, CHAP can calculate pore radius and hydrophobicity profiles. Additionally, the protein can be simulated in its native lipid bilayer environment in order to assess the behaviour of water inside the channel pore. From the resulting simulation trajectory data, CHAP can derive water density and free energy profiles that aid the identification of hydrophobic gates. Yellow arrows indicate steps of the pipeline that can be carried out on structural data alone while cyan arrows indicate those steps that are based on subsequent molecular dynamics simulation. The black arrows indicate the location of the 9’ ring of leucine residues.

The example illustrated in Figure 2 is a crystal structure of a mammalian 5-HT_3_ receptor (PDB ID: 4PIR) in which the existence of a hydrophobic gate has been demonstrated by previous studies ^21–23^ Initial structural analysis with CHAP (i.e. without simulation) yields plots of both radius and hydrophobicity for the entire channel pore. The simple radius plot shows the size of the transmembrane section of the pore to be roughly constant at ~0.3 nm radius, however, the accompanying hydrophobicity plot also reveals this section to be somewhat hydrophobic, especially in the region of the 9’ ring of side chains.

To test for the presence of a hydrophobic gate within this region, the structure was inserted in a lipid bilayer system ready for equilibrium MD simulations using GROMACS. Briefly, the starting structure was embedded in a POPC bilayer using a serial multiscale procedure ^24^ and the resulting protein-bilayer system was then simulated using the OPLS united atom force field ^25^ with united-atom lipids ^26^ and the TIP4P water model ^27^ The C*α* atoms of all protein residues were harmonically restrained at their positions in the experimentally determined structure (see **Appendix A** for details of the simulation parameters). These restraints prevent any gating transitions or other large-scale conformational transitions, whilst still permitting e.g. side chain reorientation. This ensures that any functional annotation produced by CHAP is as relevant as possible to the input structure itself, whilst simultaneously avoiding an unphysically rigid pore surface.

As shown in **Figure 2**, this allows calculation of the water density within the pore together with a free energy profile for hydration of the pore. The time-dependent behaviour of water during this particular simulation is also displayed in **Figure 3**. This shows that although the pore is fully hydrated in the initial configuration of the simulation system, water is expelled almost immediately (within < 0.5 ns), leaving a 1 nm long region of the conduction pathway fully dehydrated. Consistent with our previous studies^10^, this rapid de-wetting was also observed within the first few nanoseconds in three independent MD simulations, each of which was run for a total of 100 ns. However, the rapid (< 10 ns) nature of this dewetting indicates that much shorter simulation times are likely to be sufficient, and in this case 3x 30 ns repeats would reveal a similar result. As discussed above, it is important to note that CHAP is designed to annotate specific experimentally determined structural states i.e. it is not intended to simulate major conformational rearrangements. Thus relatively short simulation times are likely to be sufficient. Using this approach and the profiles calculated by CHAP, it becomes immediately apparent that the transmembrane pore of this particular conformation of the 5-HT_3_ receptor contains a hydrophobic gate (**Figure 2**). This would not have been revealed if only a radius profile such as that shown in **Figure 1A** had been considered.

**Figure 3:**
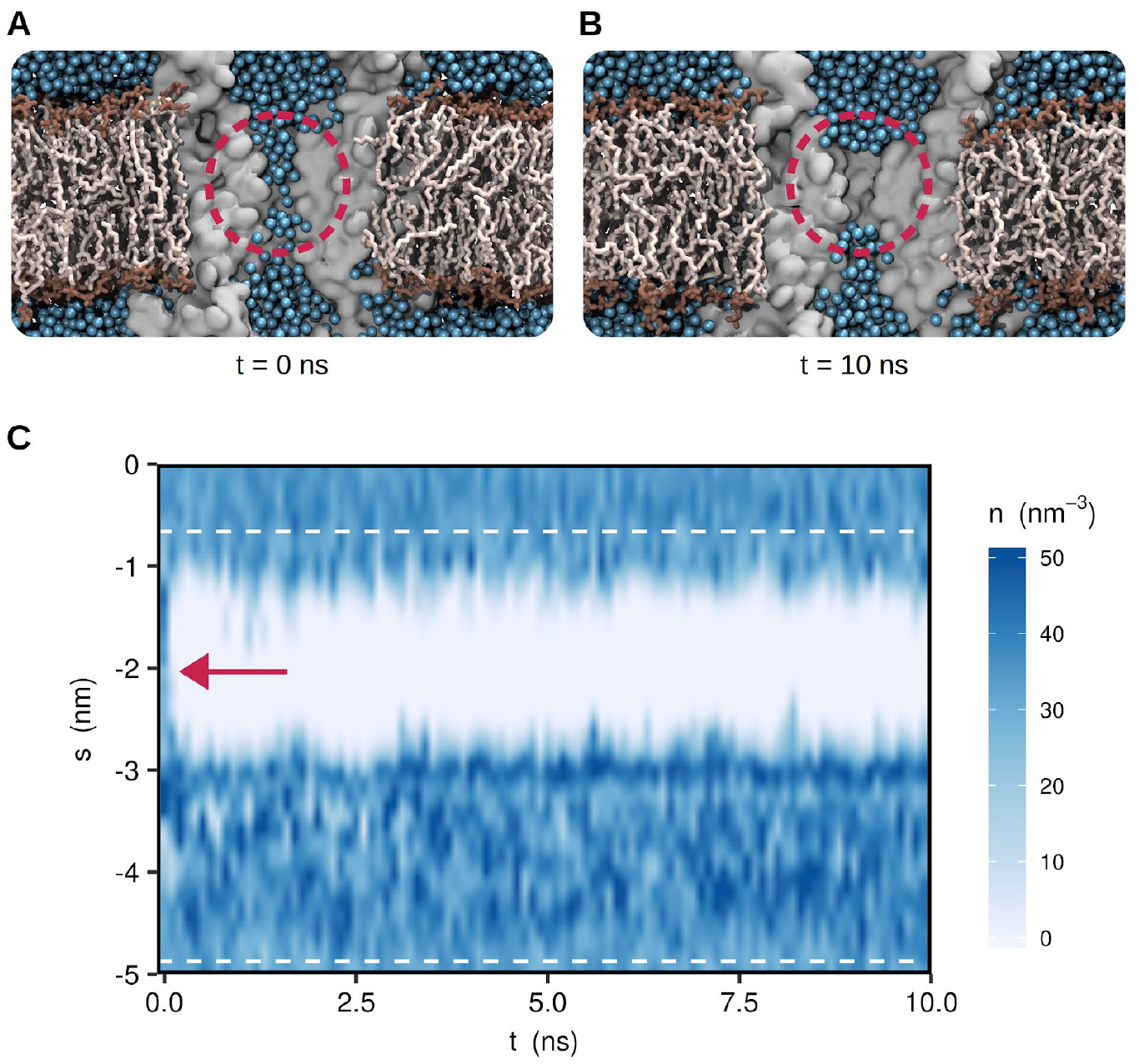
Hydrophobic gates can be identified through equilibrium MD simulation. (A) Cross-section of the starting structure for MD simulation of a 5-HT_3_ receptor (PDB ID: 4PIR) embedded in a POPC lipid bilayer. The protein is represented by a grey surface with two of five subunits omitted for visual clarity. Only the transmembrane (TM) domain is shown. Water oxygens are represented by blue spheres and lipid molecules are shown in liquorice representation. Note that the entire channel pore is initially filled with water. (B) Snapshot from the MD trajectory clearly exhibiting a de-wetted region (highlighted with red circle) in the TM domain of the channel. Note that the channel pore is not physically occluded. (C) Water density profile in the receptor pore over time. An approximately 2 nm long region of the TM domain indicated by a red arrow de-wets at the beginning of the simulation and remains devoid of water throughout.

In addition to the examples described above, we have also applied CHAP to a recently solved structure of TRPV4, a member of the TRP family of cation channels where hydrophobic gating has also recently been reported^28^. CHAP analysis of this structure reveals that although the lower gate may be able to pass hydrated Na^+^/K^+^ ions, this region is also highly hydrophobic thus presenting a major energetic barrier to hydration (**Figure S1**). This supports previous reports of hydrophobic gating in TRP channels^29,30^ and suggests that further investigation of this effect in TRPV4 is warranted.

## Implementation of CHAP

Thus although tools that calculate pore radius alone have proven exceptionally useful it is clear that additional information is required if more accurate predictions are to be made. However, a simple extension of the popular HOLE program with added functionality was not practical for several reasons. In particular, HOLE was originally developed for the analysis of static protein structures and is unable to read common trajectory data formats or process time-dependent structures similar to those generated by MD simulation. We therefore decided to implement this new channel annotation package as a completely new and independent tool: CHAP is written in C++ and is based on the trajectory analysis framework of the popular MD simulation software GROMACS ^31^.

The algorithm used by CHAP to calculate these different channel profiles is outlined in **Figure 4**. Starting from an input structure (either an experimentally derived PDB file or a single frame from a MD trajectory), the first step is to define the physical dimensions of the pore. A probe-based method is employed, similar to that used by HOLE, resulting in a sequence of discrete probe positions and associated pore radii. A continuous spatial curve is then determined from these probe positions by means of B-spline interpolation. A continuous radius profile is then calculated by interpolating the probe radii along this curve (**Figure 5**).

**Figure 4:**
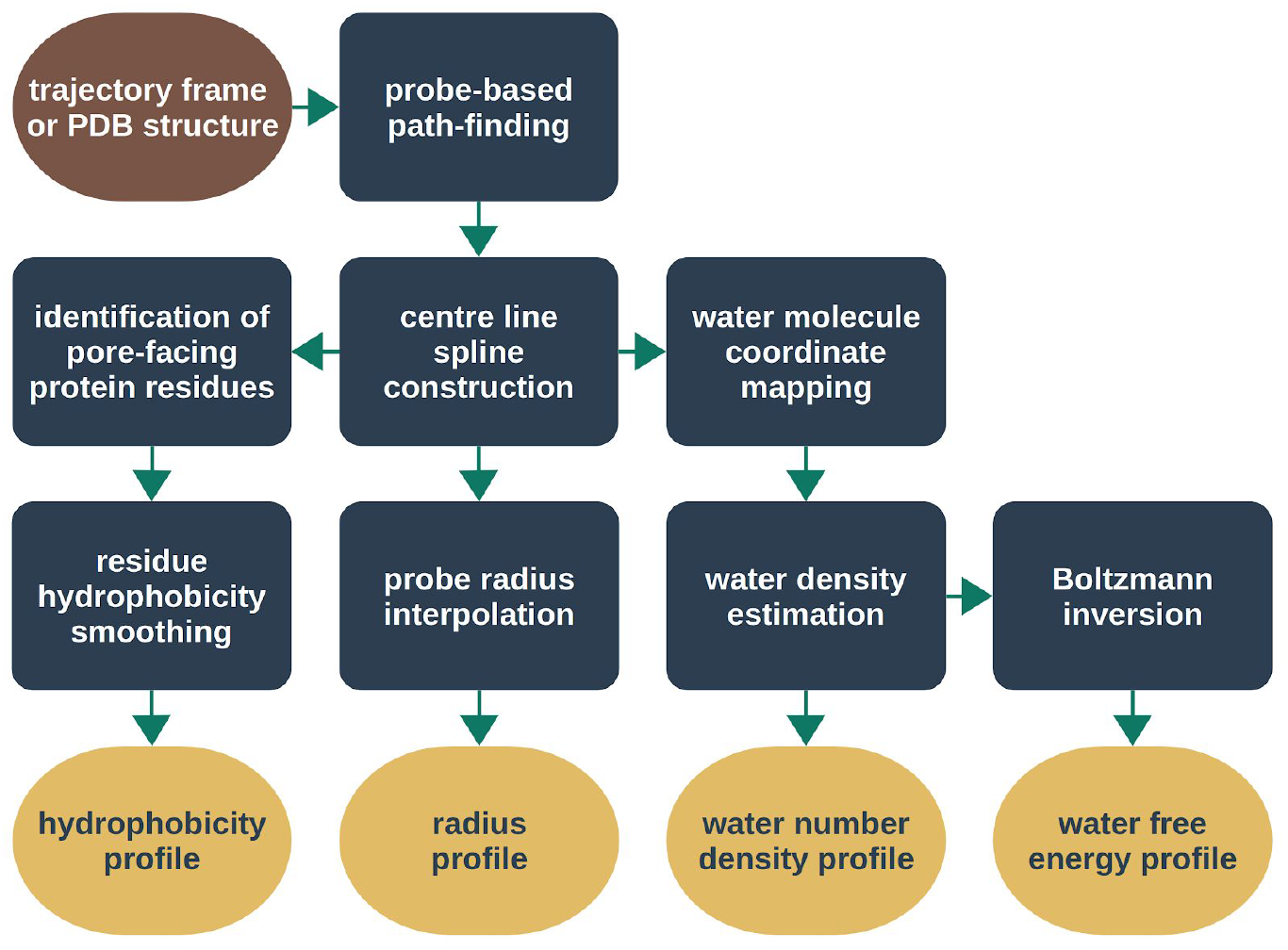
Workflow diagram of data processing in CHAP. For an individual PDB structure this procedure is carried out once. For trajectory data, the above procedure is carried out individually for each frame, i.e. for each structural configuration at a sampled time point. The resulting hydrophobicity, radius, density, and free energy profiles are averaged over all time steps and confidence bands are estimated from the standard deviation over time.

This interpolating spline curve represents the centre line of the pore and defines a coordinate system that forms the basis for all subsequent calculations. In this channel coordinate system, the position of a particle (e.g. a water molecule or a protein side chain) is represented by its distance to the closest point on the centre line and the distance along the centre line between this point and a reference point (**Figure 6A**). By convention, the reference point is the projection of the centre of geometry of the overall channel protein onto the centre line.

Next, a hydrophobicity profile is determined for the residues that line the pore. By converting residue positions to channel coordinates, CHAP identifies all residues within a cutoff distance from the pore surface and then selects only those whose side chains point inwards towards the pore. The resulting discrete values for hydrophobicity at the positions of the individual residues are then converted into a continuous hydrophobicity profile along the centre line by means of kernel smoothing ^32^.

If the input structure contains water molecules, e.g. from an MD simulation, then CHAP will also estimate the solvent density along the centre line. To achieve this, water molecule positions are first transformed into channel coordinates, and then a one-dimensional probability density is calculated by kernel density estimation ^33^. Together with the radius profile, this allows us to calculate the number density of water molecules throughout the channel structure. It is from this value that the related water free energy profile can then be estimated using Boltzmann inversion.

When the input to CHAP is an MD trajectory, the above procedure is repeated independently for each frame. CHAP will then calculate summary statistics (mean, standard deviation, minimum, and maximum) over time for each point in the channel profiles. The following sections now describe these different analysis steps in more detail.

## Pore Geometry and Centre Line

The path-finding algorithm used by CHAP is based on a hard sphere representation of the protein, in which each atom is associated with a sphere of radius *r_i_* centred at the position of the atom, **Q**_*i*_. **Figure 5A** illustrates how a pathway is found by squeezing a spherical probe through this collection of van der Waals spheres. Starting from an initial position inside the channel pore, **P**_0_, the probe is moved in steps of Δ*υ* along the direction of a unit vector **e**_*υ*_ pointing in the overall direction of the permeation pathway:

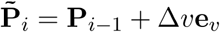

**Figure 5:**
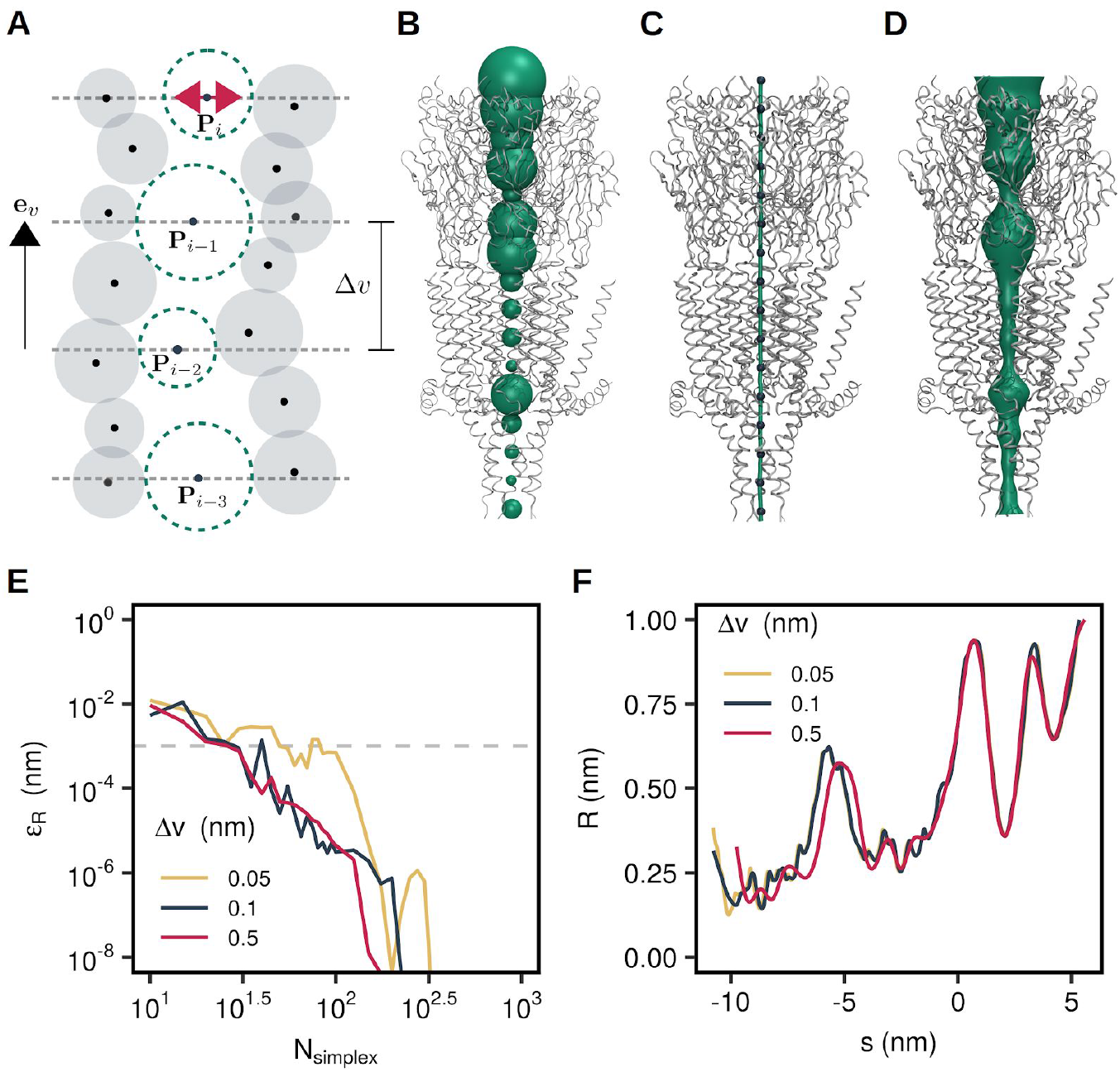
Pore geometry calculations in CHAP. (A) Schematic representation of probe-based pathway finding. The position of a spherical probe is optimised in subsequent parallel planes to maximise its radius without overlapping the van der Waals sphere of any ion channel atom. (B) Configuration space view of probe positions (green spheres) in the permeation pathway of a 5-HT_3_ receptor structure (PDB ID: 4PIR). For visual clarity a large probe step of Δ*ν* = 1 nm was used here. (C) The probe centre positions (blue spheres) are interpolated with cubic B-splines to yield a continuous spatial curve (green tube) representing the pore centre line. (D) A surface representation of the channel pore is generated by extruding a circular cross-section along the centre line spline curve. (E) Convergence of the Nelder-Mead algorithm carried out to optimise the probe position in each parallel plane. The radius error falls below a threshold of 0.001 nm within 100 optimisation steps irrespective of the value of the probe step. The reference radius profile is calculated with a very large number of 2000 Nelder-Mead iterations. (F) Influence of probe step on radius profile resolution. While a probe step of 0.5 nm only captures the general feature of the radius profile, decreasing the probe step from 0.1 nm to 0.05 nm leads only to minimal changes.

The resulting point is then used to initialise a numerical optimisation procedure seeking to maximise the radius of the probe,

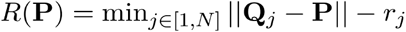

without overlapping the van der Waals sphere of any protein atom. This optimisation procedure is carried out in the plane supported by 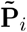 and spanned by the unit vectors **e**_*u*_ and **e**_*w*_ which are chosen to be perpendicular to both **e**_*υ*_ and one another. The optimal probe position

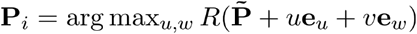

is accepted as a point on the pore centre line and a further step in the direction of **e**_*υ*_ is carried out. This procedure is repeated until the probe radius exceeds a threshold, at which point it is assumed that the probe has left the pore. Subsequently, the iteration is carried out a second time moving in the direction of −**e**_*υ*_ and starting from point **P**_1_.

In the original algorithm used by HOLE the optimisation of the probe position was performed by means of simulated annealing ^5^. However, being a global optimisation procedure, simulated annealing is prone to allowing the probe to jump out of the protein pore. The downhill simplex algorithm ^34^ was therefore implemented in CHAP alongside the simulated annealing procedure and this latter method was found to be generally more robust.

This procedure has three free parameters: The probe step size, Δ*υ*, the number of Nelder-Mead iterations, *N*_simplex_, and the initial probe position, *P*_0_. For a sufficiently symmetric protein with a central pore, *P*_0_ can simply be set to the centre of geometry of the overall channel protein. The step size needs to be small enough to resolve all structural features of interest. As illustrated in **Figure 5F**, a probe step of 0.1 nm (the van der Waals radius of a hydrogen atom) is sufficient and the radius profile does not change significantly when the probe step is decreased further. Figure 5E shows the dependence of the error in the radius profile on the number of simplex iterations and reveals that 100 steps are sufficient to converge the profile to within a tolerance of 0.001 nm even for a very small probe step. This is the case not just for this particular 5-HT_3_ receptor structure, but also for a wide range of other ion channel proteins (**Figure S2**).

The points **P**_*i*_ on the centre line are then connected into a continuous spatial curve **S**(*s*) by means of B-spline interpolation as detailed in **Appendix B** so that the curve passes exactly through all centre line points:

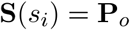

A corresponding continuous radius profile is then found by interpolating the probe radii along the centre line curve. The main advantage of a continuous centre line curve over individual points is that the curve can be used to define a simple one-dimensional reaction coordinate for the motion of particles through the channel pore, as is detailed in the following section. Additionally, it can be used to generate a pore surface representation through the path extrusion technique as shown in **Figure 5D**. Importantly, for radially symmetrical pores the radius profiles calculated using CHAP are almost identical to those calculated using HOLE (**Figure S3**). However, it should be noted that the S coordinate calculated by CHAP follows the pathway of the pore, and so for non-linear pathways this may differ from the Z-coordinate calculated by HOLE.

## Water Density and Free Energy

We have previously shown water density can be used to estimate a free energy profile for pore hydration and how this acts as proxy for measurement of ion permeation^21^. To achieve this, the centre line spline is used to define a curvilinear coordinate system, in which the position of each particle is described by its distance from the spline curve, *ρ*, the arc length parameter value corresponding to the closest point on the spline curve, *s*, and the angle, *ϕ*, enclosed between the vector connecting this point and the particle position and the normal vector of the spline curve (**Figure 6A**). The resulting coordinate system can be viewed as a polar version of the Frenet-Serret frame, providing a generalisation of the canonical cylindrical coordinate system, where the *z*-coordinate is replaced by the length along the arc of the centre line.

**Figure 6:**
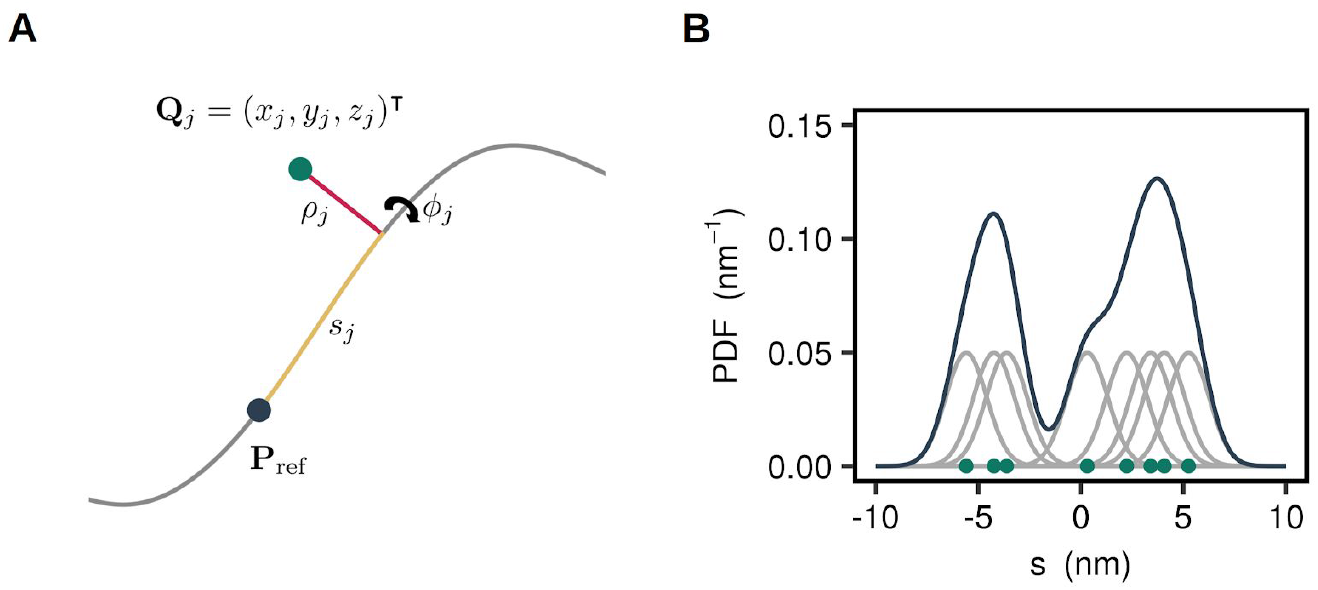
Determination of water density. (A) Cartesian coordinates of a water molecule (green) are converted into a curvilinear coordinate system by projecting its position onto the channel centre line (grey). The new coordinates are then expressed in terms of the distance along the centre line (yellow) between the projection base point and a reference point (dark blue), the distance from the water molecule to the centre line (red), and an angle around the centre line. (B) Conceptual representation of kernel density estimation: Each discrete water molecule position is associated with a Gaussian probability distribution and the sum of all Gaussians yields the probability density function of water along the channel centre line.

Unfortunately, there is no analytical expression relating the Cartesian coordinates of a particle to the curvilinear centre line coordinates and this mapping has to be accomplished numerically by finding the minimal distance between the position of a particle and the spline curve. Given the position of the *j*-th particle in Cartesian coordinates, **Q**_*j*_ = (*x_j_*, *y_j_*, *z_j_*)^⊤^ the corresponding spline coordinates are found from

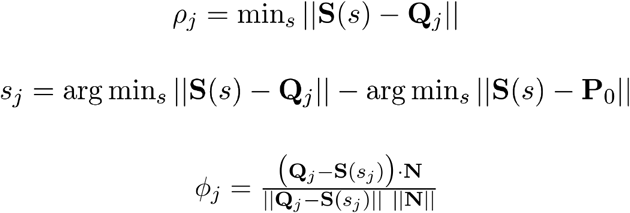

where the minimisation problem is solved using the method of ^35^. Note that in the definition of the *s*-coordinate the projection of the initial probe position is subtracted. This ensures that the offset of the *s*-coordinate is consistent across multiple trajectory frames by setting it to zero for the initial probe position (which typically corresponds to the centre of geometry of the protein).

The *s*-coordinate can be used as a one-dimensional collective variable along which to determine water density and the corresponding free energy profile. The probability density *P*(*s*, *ρ*, *ϕ*) of finding a water molecule at the given centre line coordinates is given by the Boltzmann relation,

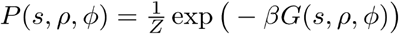

where *G*(*s*, *ρ*, *ϕ*) is the Gibbs free energy, *Z* is the partition function, and *β*^−1^ = *k*_B_*T* with *k*_B_ being the Boltzmann constant and *T* the simulation temperature. In order to obtain a one-dimensional free energy profile, i.e. *G*(*s*, *ρ*, *ϕ*) = *G*(*s*) the lateral degrees of freedom are integrated out according to

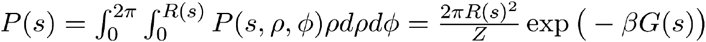

where it is assumed that the cross-sectional area of the pore is approximately circular with a local radius *R*(*s*). Inverting the above relationship yields an expression for the free energy of the water molecule,

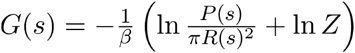

into which the partition function enters only as an additive constant. This constant is determined by the convention that the free energy in the bulk outside the protein should be zero, i.e. *G*(*s*) → 0 as *s* → ±∞.

The probability density appearing in the above expression can be estimated directly from the MD trajectory by means of kernel density estimation:

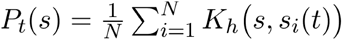

Here *s_i_*(*t*) is the position of the *i*-th water molecule at time *t*, *N* is the overall number of water molecules, and *P_t_*(*s*) denotes the probability density estimated at time *t*. A Gaussian kernel of the form

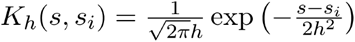

is employed here. A conceptual representation of kernel density estimation is shown in **Figure 6B**.

The bandwidth *h* is a free parameter in kernel density estimation that fulfills a function similar to the bin width in histograms and care must be taken when selecting a value for this parameter as the resulting density profile is sensitive to it. A value for this parameter of 0.24 nm has been suggested for identifying liquid interfaces in three dimensions ^36^, whereas a value of 0.1 nm has been used in a study of pore de-wetting in the bacterial GLIC channel ^16^. It is also possible to estimate an appropriate bandwidth directly from the data ^33^. This method minimises the asymptotic mean integrated square error (AMISE) of the density estimate.

**Figure 7** uses the example of the 5-HT_3_ receptor to illustrate how the water density and free energy profiles depend on the bandwidth parameter. For very small values of *h*, the water density estimate is very noisy and in low density regions it drops down to zero, so that the corresponding free energy at these points is singular. Conversely, if a very large bandwidth is chosen, the estimate of *P*(*s*) is near uniform throughout the pore and the water density estimate exhibits over-densities at points where the pore radius is particularly small. In this case the free energy profile exhibits no clear barriers or wells.

**Figure 7:**
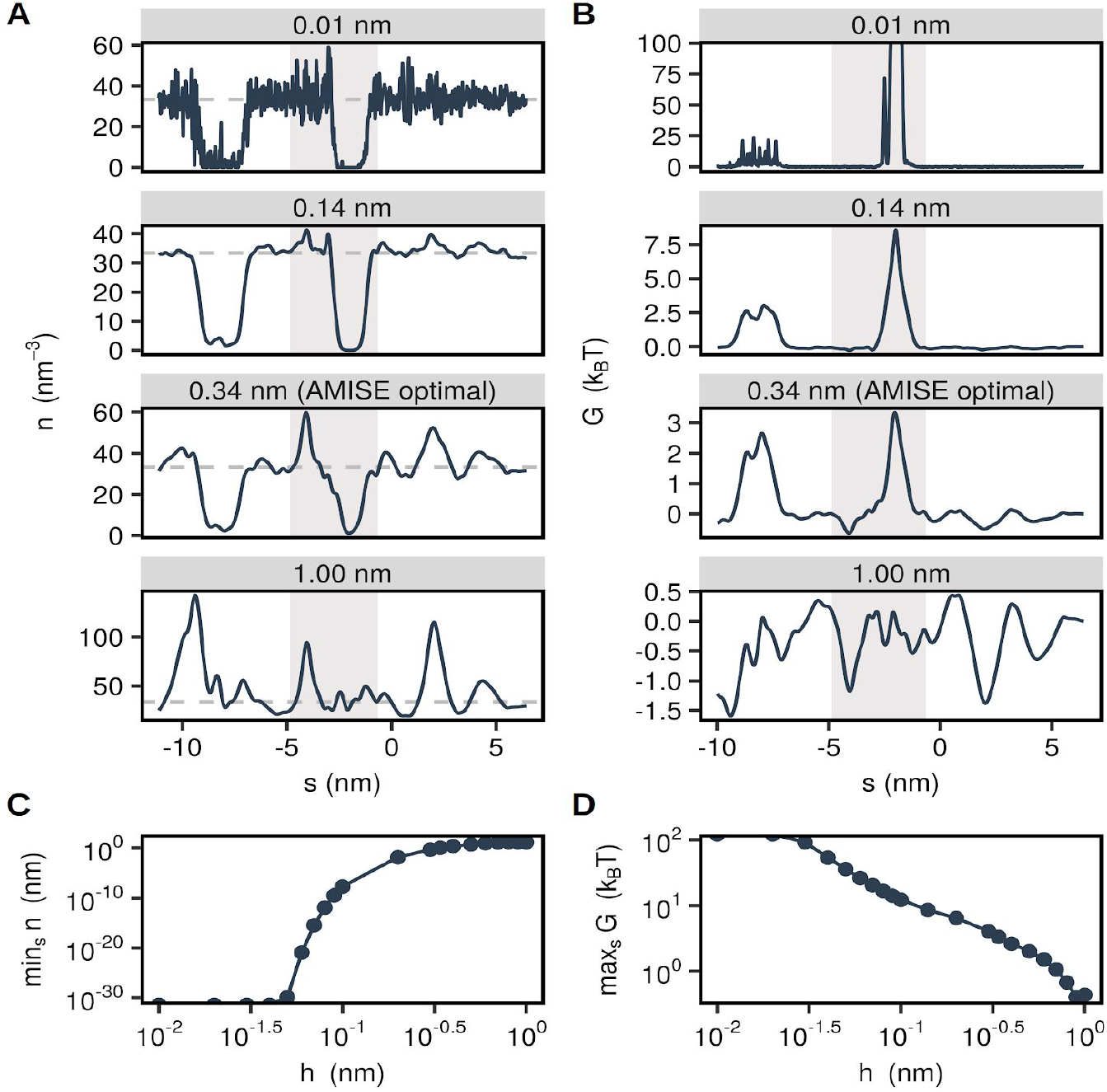
Influence of bandwidth parameter. (A) Water number density along the permeation pathway of the 5-HT_3_ receptor for various values of the kernel bandwidth. A value of 0.14 nm corresponds to the approximate size of a water molecule and a value of 0.34 nm corresponds to the statistically optimal bandwidth. (B) Corresponding free energy profiles calculated via Boltzmann inversion of the density profiles. The grey shaded area in (A) and (B) represents the transmembrane domain. (C) Minimum of number density profile in dependence of bandwidth. (D) Maximum of free energy profile in dependence of bandwidth.

The AMISE-optimal bandwidth, which in this case is 0.34 nm, finds a balance between these two extremes. The overall density profile is free of noise and the lowered water density in the region of the hydrophobic gate is readily apparent. However, there are still some regions where the water density is unphysically large. By comparison, a profile estimated using a bandwidth of 0.14 nm (the approximate radius of a water molecule) yields a water density that never much exceeds the experimental bulk density of water while still showing little sign of noise. Using a bandwidth fixed at the size of a water molecule also permits a more direct comparison between different channel structures, while the AMISE-optimal bandwidth may differ across channel structures of different overall pore volume. It is therefore recommended to use a default value of 0.14 nm.

## Hydrophobicity Profiles

In addition to calculating the channel radius and water density profiles inside the pore, it is also necessary to quantify the hydrophobicity along the conduction pathway. For cylindrical nanopores, it has been found that the free energy difference between hydrated and dehydrated pore states is given by

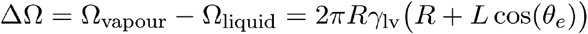

where *R* and *L* are the radius and length of the nanopore, *γ*_1v_ is the liquid-vapour surface tension of water, and *θ_e_* is the contact angle from the Young-Dupré equation ^13^. Determining a hydrophobicity profile for a channel therefore requires knowledge about the amino acid residues facing a given section of the pore and their associated contact angle. The individual hydrophobicity values at the positions of the amino acids can then be combined into a continuous hydrophobicity profile by means of kernel smoothing.

CHAP can automatically identify pore-facing residues, i.e. residues exposing their amino acid side chains to the channel pore, via a two-step procedure. First, all residues whose centre of geometry lies within a given cutoff, *R_thres_*, from the pore surface are determined. To accomplish this, the position of the centre of geometry of each residue is mapped from Cartesian coordinates to centre line spline coordinates so that each residue is associated with a distance from the centre line, 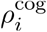, and a distance along the centre line, 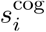, where the index runs over all protein residues. If

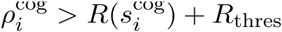

the *i*-th residue will be considered pore-lining, where 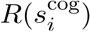 is the pore radius at the position of the residue. A typical threshold value for an *α*-helical pore is 0.75 nm.

Second, for each pore-lining residue, the position of the *α*-carbon is also mapped to pathway coordinates 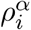 and 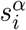. If the distance from the *α*-carbon to the pore centre line is larger than the distance between the centre of geometry of the overall residue and the pore centre line, i.e. if

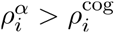

then the *i*-th residue will be considered pore-facing. The property of being pore-lining and pore-facing is encoded in two binary indicator variables, whose values are written to the temperature factor and occupancy fields of a PDB file of the input structure to allow easy visualisation of these residues. If CHAP is run with a trajectory as input, the above analysis is performed repeatedly (once for each frame) and the values are written to the PDB file correspond to the time average of the indicator variables. **Figure 8A** illustrates the distinction between pore-lining and pore-facing residues in the 5-HT_3_ receptor.

**Figure 8:**
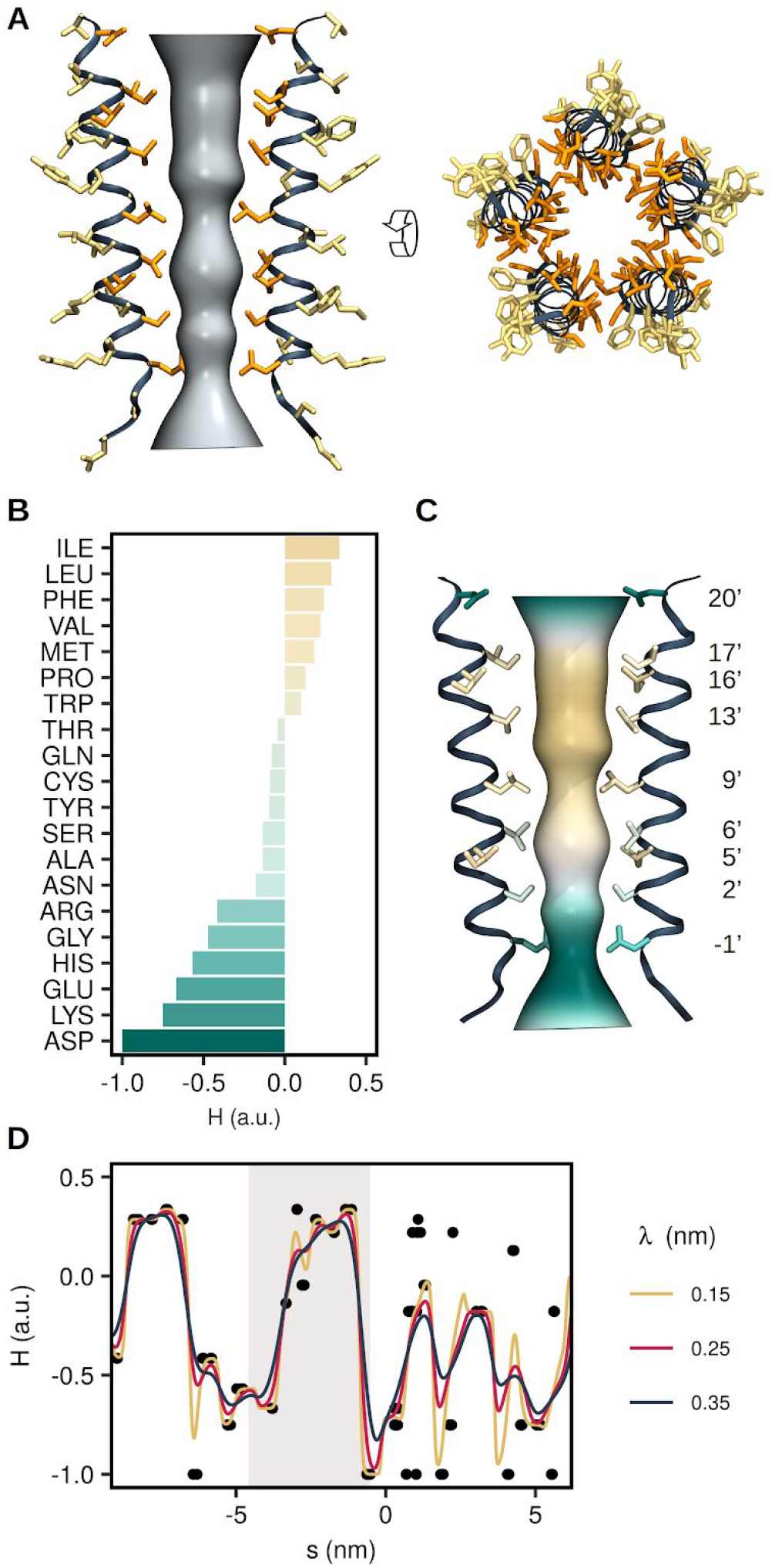
Hydrophobicity profile of pore-lining residues. (A) Classification of pore-lining residues in the M2 helix bundle of the 5-HT_3_ receptor transmembrane domain into side chains pointing away from the pore (yellow) and facing the pore (orange). For visual clarity only two out of five helices are shown in the side view. (B) Amino acid hydrophobicities according to the scale proposed by Wimley and White. In CHAP the original hydrophobicities are rescaled into the interval from −1 (very hydrophilic) to +1 (very hydrophobic) to facilitate an easy comparison between different hydrophobicity scales. (C) Permeation pathway coloured according to the Wimley-White hydrophobicity of the pore-facing residues. (D) Hydrophobicity profile of the entire 5-HT_3_ receptor pore. Black circles represent the position and hydrophobicity of the pore-facing residues (averaged over all five subunits). Lines represent the kernel-smoothed continuous hydrophobicity profile underlying the surface colouring in panel (C). Different values of the smoothing span, λ, influence the exact magnitude of hydrophobicity fluctuations, but the overall distinction between hydrophobic and hydrophilic regions along of the pore remains unaffected. The shaded area corresponds to the transmembrane region shown in panels (A) and (C).

To the best of our knowledge no experimental data quantifying the contact angle at the interface between water and specific amino acids are available. However, various amino acid hydrophobicity scales have been proposed in the literature. An early hydropathy scale based on a collection of experimental observations has been suggested by ^37^, whilst another has been based on measurements of the stability of *α*-helices ^38^. Other scales have used the free energy of partitioning of oligopeptides between water and a lipid environment ^39,40^, while some report free energies of transferring amino acids from water into a lipid bilayer ^41^. A biological scale has also been proposed based on the recognition of *α*-helices by the endoplasmic reticulum translocon ^42^, whilst another is derived from MD simulations of a water nanodroplet on a planar layer of amino acids ^43^.

For comparability, the hydrophobicity scales listed above have been shifted and scaled to the interval ranging from −1 for the most hydrophilic to +1 for the most hydrophobic residues while preserving their natural zero (**Figure S4**). Note that this contrasts with the physical definition of hydrophobicity in terms of cos*θ*_*e*_ and instead follows the biochemical convention of associating a larger value with a more hydrophobic residue. While the exact hydrophobicity value assigned to each amino acid differs to some degree across scales (**Figure S4**), we found no marked difference in their ability to predict pore de-wetting in a recent analysis of nearly 200 ion channel structures ^44^. By default, CHAP employs the Wimley and White scale, but users are free to select a alternative scales that may be more appropriate for their structure.

CHAP uses the normalised hydrophobicity scales to associate a hydrophobicity value, *H_i_*, with each residue position 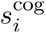. A continuous hydrophobicity profile is then calculated from these discrete hydrophobicity values using a Nadaraya-Watson kernel-weighted average. Mathematically this is expressed as

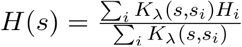

where

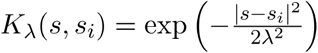

is a Gaussian kernel. The smoothing span λ is chosen to reflect the finite extension of an amino acid residue. Since the pitch of an *α*-helix is approximately 0.15 nm per residue and on average every third residue is pore-facing, a value of ~0.225 nm (for a width of the Gaussian kernel of 0.45 nm) can be considered reasonable. Note that far away from any residues (e.g. in the bulk water region outside the channel pore) *H*(*s*) will decay exponentially to zero.

In **Figure 8D** this is illustrated on the 5-HT_3_ receptor using the Wimley and White hydrophobicity scale. As can be seen, there is a considerable increase in hydrophobicity in the region of the transmembrane domain containing the hydrophobic gate. This section of the profile is shown in **Figure 8C** mapped onto a 3D representation of the pore together with the M2 helices of the 5-HT_3_ receptor, illustrating how the hydrophobicity of the pore-facing residues influences the hydrophobicity profile.

## Conclusion

In this study we present a versatile computational tool for the functional annotation of ion channel structures: the Channel Annotation Package CHAP. This tool implements a methodology for predicting the conductive state of ion channel structures that goes beyond a measurement of the pore radius to include profiles of pore hydrophobicity as well as of water density and free energy based on MD simulations of water behaviour within and experimentally determined channel structure. Crucially, this enables CHAP to identify hydrophobic gates which cannot be detected using methods based purely on measuring the physical dimensions of the pore alone.

Importantly, the analysis performed by CHAP is computationally inexpensive compared to more complex simulation techniques such as computational electrophysiology ^45^ and is therefore highly suitable for the high-throughput analysis of the rapidly growing number of ion channel structures. The utility of this approach is demonstrated in a recent study ^44^ employing CHAP to investigate the prevalence of hydrophobic gates in ~200 different ion channel structures.

CHAP is made available as free and open-source software. Copies of the source code can be obtained from www.channotation.org and include scripts for visualising results in the molecular graphics programmes VMD ^46^ and PyMOL ^47^.

## Acknowledgements

This work was funded by grants from the BBSRC, the EPSRC, and the Wellcome Trust. GK holds a studentship funded by the EPSRC.

## Appendix A: Molecular Dynamics Simulation Protocol

Equilibrium molecular dynamics simulations of the 5-HT_3_ receptor were performed using GROMACS 2018. Starting structures were generated from the original crystal structure (PDB ID: 4PIR) with missing atoms being added by the WHAT IF tool ^48^. This starting structure was embedded in a DOPC bilayer using an established serial multiscale protocol 24. The resulting protein-bilayer system was converted back to an atomistic representation and solvated in 150 mM NaCl solution. The CHARMM36 all-atom force field^49^ was used together with the TIP3P model of water ^27^. Simulations of the TRPV4 channel (PDB ID: 6BBJ) followed the same protocol.

Long-range electrostatic interactions are treated using the particle mesh Ewald (PME) method ^48,50^ employing a short range cutoff of 1 nm and a Fourier spacing of 0.12 nm. To permit subsequent determination of the Gibbs free energy from the water molecule distribution, the system is simulated in the isothermal-isobaric (NPT) ensemble at a temperature of 310 K and a pressure of 1 bar. A v-rescale thermostat ^51^ with a coupling constant of 0.1 ps is used for temperature control and a pressure is maintained semi-isotropically using the method of ^52^ with a coupling constant of 1 ps. The canonical leapfrog method with a time step of 2 fs is used to integrate three independent copies of the system for 100 ns with bonds constrained through the linear constraints solver algorithm LINCS ^53^. Additionally, the C*α* atoms of all protein residues are placed under a harmonic restraint with a force constant of 1000 kJ/mol/nm^2^ to ensure that the configuration of the simulated channel does not deviate from the experimentally determined structure. A simulation length of 100 ns likely exceeds that required, since the largest permitted protein motions are side chain reorientations, which typically occur on a time scale of ~10 ns and most pore dewetting events appear to occur even more rapidly. For the examples described in this study, analysis of 3x 30 ns simulations would produce similar results.

## Appendix B: B-Spline Curves

Formally, a spline curve of degree *p* along some parameter *s* can be written as the linear combination

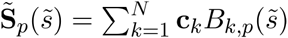

where

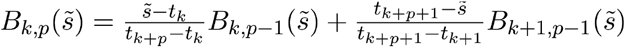

with 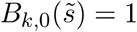 if 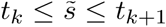 and 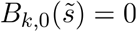 otherwise are the recursively defined B-spline basis functions. Here 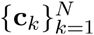 denotes the spline curve control points and 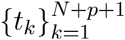 is referred to as knot vector. Note that a zero division in the above equations is treated as 0/0 = 0.

In order to find the spline curve that interpolates the centre line points **P**_*i*_ from the pore finding algorithm, one demands

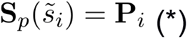

which requires a parameterisation of the discrete point set. Ideally, this should be the distance between the points along the arc of the spline curve, but unfortunately this distance is not known *a priori*. Instead, the commonly applied chord length approximation is used here, in which points are parameterised by their Euclidean distance according to:

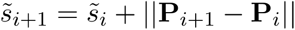

Importantly, it has been also shown that this choice of parameterisation guarantees full approximation order for spline curves up to third degree ^54^.

For a cubic spline curve 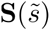, where the subscript *p* = 3 has been dropped for clarity, Equation (*) together with the knot vector

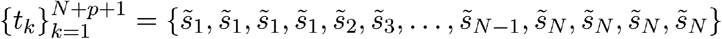

and hermite boundary conditions then leads to a tridiagonal linear system, from which the control points **c**_*k*_ can be determined efficiently through the Thomas algorithm. The choice of cubic splines is motivated by the fact that cubic splines are twice continuously differentiable so that the curvature of the resulting spline curve is guaranteed to be continuous. At the same time, cubic splines are known to minimise overall curvature, preventing undue variation of the interpolant between the given centre line points.

The resulting spline curve is subsequently re-parameterised by arc length using a three-step procedure proposed by ^55^. In a first step, the arc length in each interval is computed according to

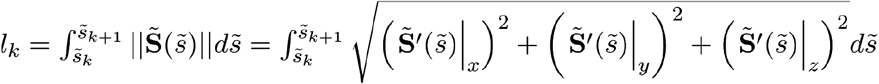

from which the overall length of the spline curve between the two openings of the pore can be calculated as:

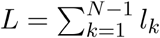

The integral in the expression for *l_k_* is solved numerically through sixth-order accurate Newton-Cotes quadrature (Boole’s rule).

The second step comprises calculating the chord length parameter values 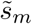 corresponding to the equidistant arc length parameter values *s_m_* = *m*Δ*s* = *mL*/*M* with *m* ∈ [0, *M*] This is equivalent to finding the root of

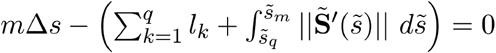

where *q* ∈ [1, *N*] is the index of the spline interval for which

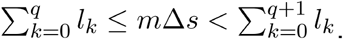

The root-finding problem is solved through the algorithm of ^56^, which is known to be the asymptotically most efficient root-finding method.

In the final step, the interpolation problem

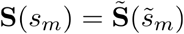

is solved to obtain a new spline curve, **S**(**s**), that approximates the original curve, but is parametrized by arc length rather than chord length. This curve is then used to describe the centre line of the channel pore.

---

**Figure S1:**
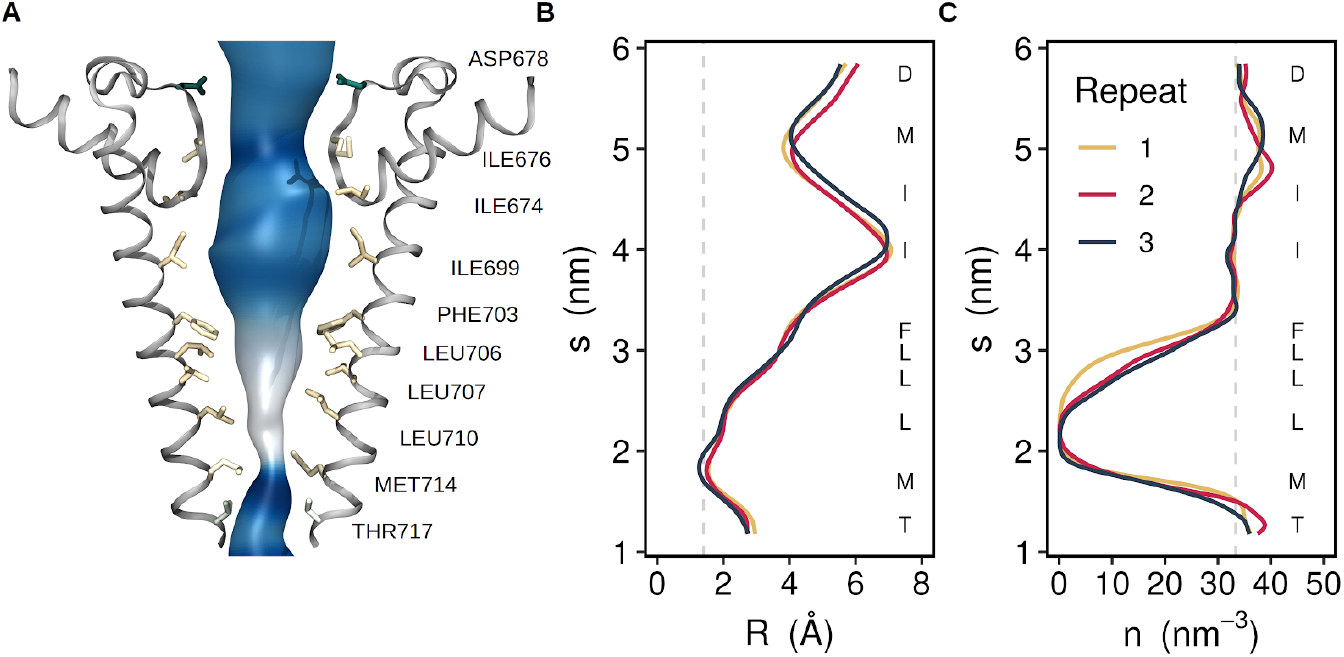
A hydrophobic gate in TRPV4. (A) Permeation pathway through the transmembrane domain of the TRPV4 structure 6BBJ. Pore-facing residues are shown in licorice representation and are coloured according to their hydrophobicity. (B) Time-averaged radius profile of the permeation pathway determined from three independent MD simulations. (C) Corresponding water number density profiles.

**Figure S2:**
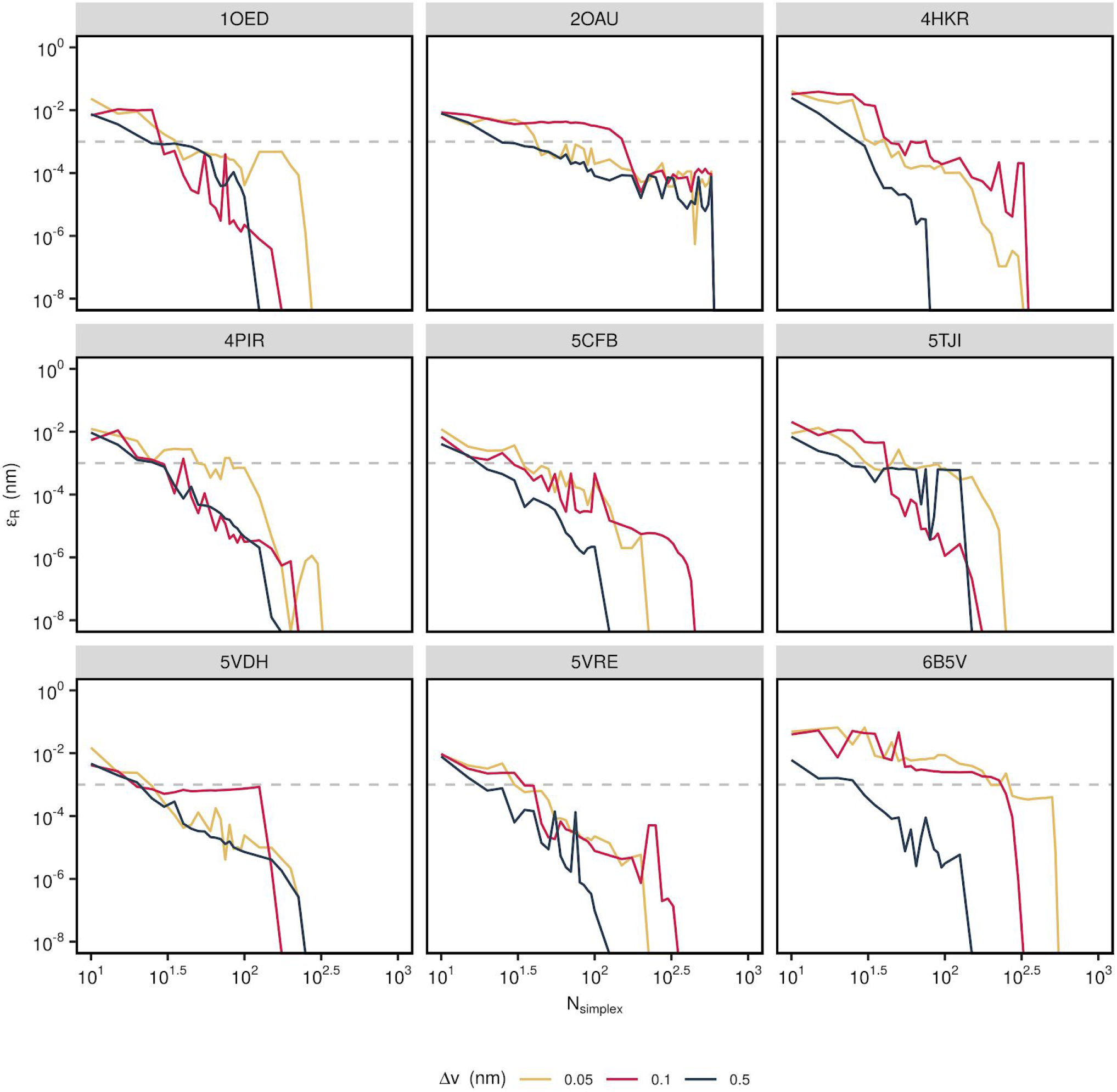
Convergence of the Nelder-Mead algorithm. The error of the radius profile calculated for nine different channel structures using three different values for the probe step is shown. Convergence to within 0.001 nm is typically reached within 100 iterations.

**Figure S3:**
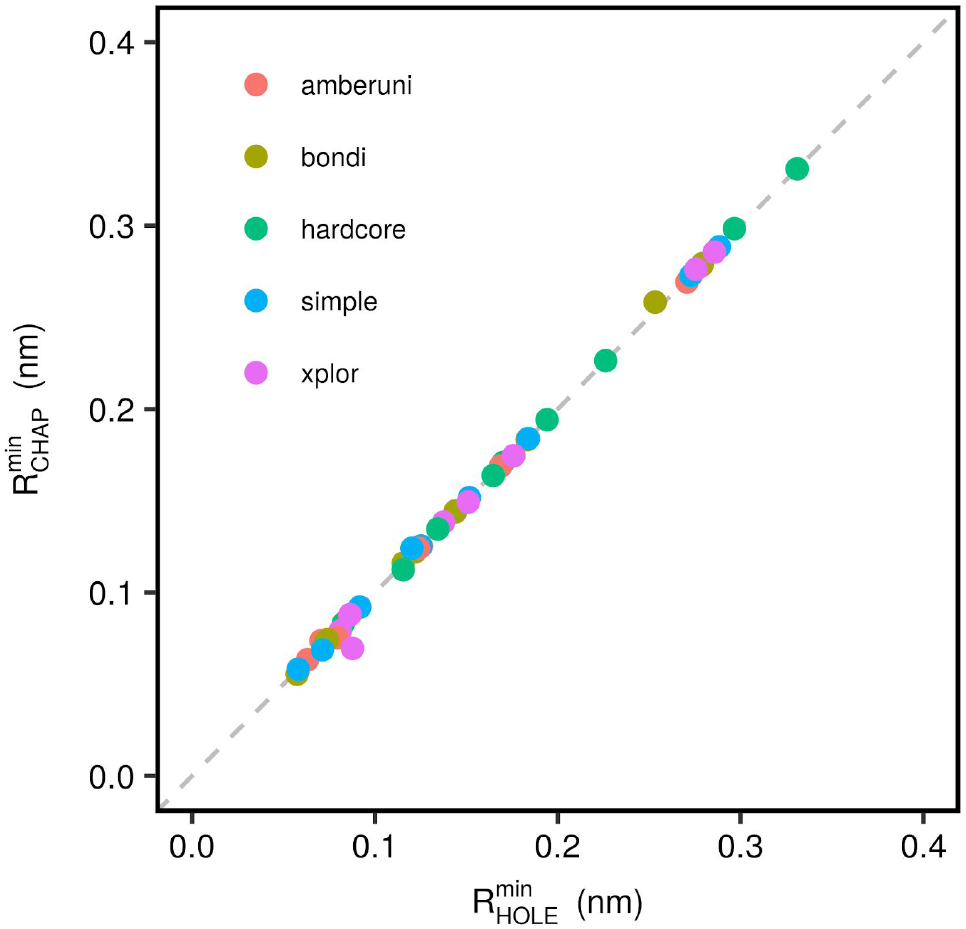
Comparison of the radius at the narrowest constriction between CHAP and HOLE. Calculated for nine representative ion channel structures using the five different van der Waals radius datasets available in HOLE. In all cases the radii agree to within 0.01 nm.

**Figure S4:**
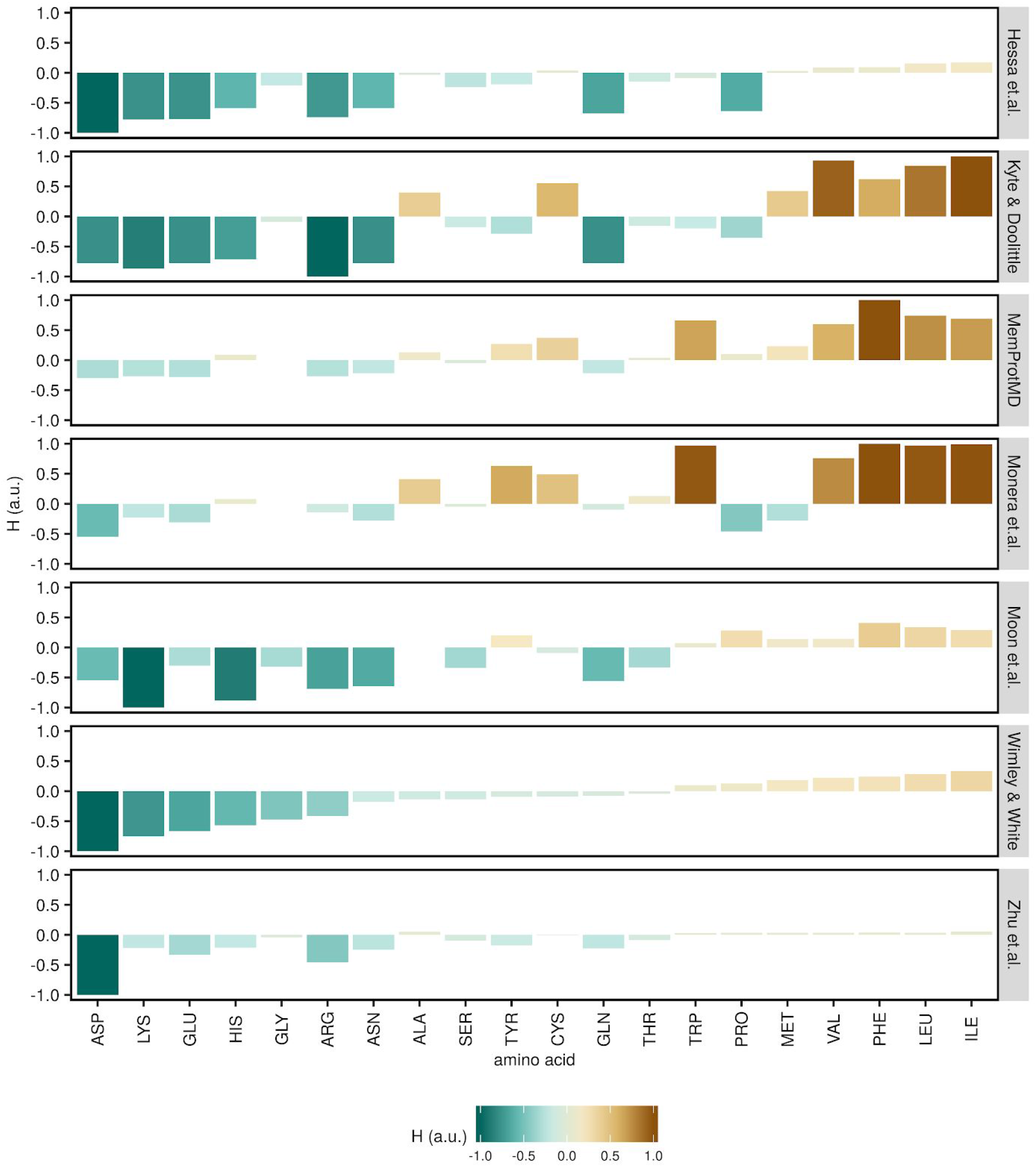
Comparison of amino acid hydrophobicity scales available in CHAP. The amino acids are listed in order of increasing hydrophobicity according to the Wimley-White scale, which is the default scale used by CHAP.

